# Decomposition trajectories in mixed-species leaf litter in cool-temperate deciduous forest with understory dwarf bamboo

**DOI:** 10.64898/2026.10.13.758829

**Authors:** Shinpei Yoshitake, Chiyuki Asai, Toshiyuki Ohtsuka

## Abstract

Litter decomposition plays a central role in nutrient cycling in forest ecosystems, but the effects of species mixing on decomposition of individual litter species remain poorly understood. We conducted a two-year litterbag experiment in a cool-temperate deciduous forest in northern Japan using litter from two dominant canopy tree species (oak and birch) and the dominant understory bamboo (Sasa). Single-species and mixed-species litterbags were used to examine litter mass loss, litter chemistry (carbon, nitrogen, and lignin), and microbial communities assessed by phospholipid fatty acid (PLFA) analysis. Oak and birch litter decomposed significantly faster than Sasa litter, whereas decomposition rate constants did not differ between the two deciduous tree species. Changes in litter chemistry relative to mass loss were similar between oak and birch but differed markedly in Sasa, indicating a distinct decomposition trajectory of Sasa. Total microbial biomass increased rapidly during the early stage of decomposition in all litter types owing to fungal colonization, whereas the fungal-to-bacterial PLFA ratio was substantially lower in Sasa litter than in deciduous tree litter. Species mixing had no significant overall non-additive effect on decomposition, although mixtures of oak and birch tended to accelerate decomposition, whereas mixtures including Sasa tended to suppress it. These changes in decomposition rate were accompanied by corresponding changes in microbial biomass, while decomposition trajectories remained largely unchanged within each litter species. Our results suggest that Sasa contributes a distinct decomposition process in cool-temperate forests and that species mixing primarily modifies decomposition rates rather than the sequence of chemical transformations during decomposition.

## Introduction

Litterfall represents the largest flux of carbon from vegetation to soil in forest ecosystems (e.g., Currie 2003; Ohtsuka et al. 2007). After entering the forest floor, most of the carbon (C) in litter is returned to the atmosphere as CO_2_ through decomposition, whereas nutrients such as nitrogen (N) and phosphorus (P) are mineralized and recycled into the soil, where they become available for plant uptake. In deciduous broad-leaved forests, where litterfall occurs synchronously each autumn, annual litter production can even exceed woody net primary production (Ohtsuka et al. 2005). Consequently, litter decomposition plays a central role in regulating nutrient cycling and biogeochemical processes in forest soils.

Numerous litterbag studies have demonstrated that litter mass decreases exponentially over time, and the rate of decomposition has commonly been quantified by the decomposition rate constant (*k*) (e.g., Olson 1963; Harmon et al. 2009). Litter decomposition is generally considered to be regulated by three major factors: the physical environment, decomposer organisms, and litter quality (e.g., Allison et al. 2006). From the perspective of energy flow, more than 90% of litter decomposition on the forest floor is carried out by microorganisms (Berg and McClaugherty 2004). Because these heterotrophic microorganisms obtain C and nutrients required for growth by decomposing litter, their activity is strongly influenced by litter chemical properties, including C, N, and lignin contents. Accordingly, the C/N ratio and lignin/N ratio are widely recognized as useful proxies of the initial decomposition rate of plant litter (Melillo et al. 1982; Talbot and Treseder 2012).

However, litter chemistry itself changes continuously during decomposition through processes such as nitrogen immobilization, nitrogen mineralization, and the accumulation of recalcitrant compounds (e.g., Blair et al. 1990; Osono and Takeda 2005; Osono et al. 2014). Likewise, microbial communities associated with decomposing litter also undergo succession as litter chemistry changes during decomposition (e.g., Berg and McClaugherty 2004; Osono 2007). Thus, litter decomposition is a dynamic process characterized not only by progressive mass loss but also by continuous changes in litter chemistry and microbial communities. Therefore, evaluating decomposition solely based on the decomposition rate may overlook the sequence of chemical and microbial changes that accompany decomposition. Evaluating this sequence, hereafter referred to as the decomposition trajectory, can provide a more comprehensive understanding of litter decomposition.

In forest ecosystems, leaf litter derived from multiple plant species usually decomposes as mixed-species litter. If litter species decompose independently, decomposition of a litter mixture is expected to equal the weighted average of the component species (i.e., an additive effect). However, the pioneering review by Gartner and Cardon (2004) and subsequent meta-analyses (e.g., Liu et al. 2020; Kou et al. 2020; Bai et al. 2026) have shown that decomposition of litter mixtures often deviates from this additive expectation. Mixture decomposition may be accelerated relative to the expected value (a synergistic effect) or suppressed (an antagonistic effect). The existence of such non-additive effects suggests that plant diversity can influence decomposition rates and thereby affect nutrient cycling, plant productivity, and ecosystem functioning. In contrast, a recent meta-analysis by Porre et al. (2020) concluded that litter decomposition is largely predictable from additive expectations, indicating that no consensus has yet been reached regarding the generality of non-additive litter-mixture effects.

Since the early 2000s, numerous studies have investigated the mechanisms underlying litter-mixture effects. Among the mechanisms proposed to explain non-additive effects, particularly synergistic effects, two hypotheses have received the greatest support. The first is resource complementarity, whereby nutrients released from high-quality litter facilitate the decomposition of lower-quality litter (e.g., Schimel and Hättenschwiler 2007; Yin and Koide 2019; Liu et al. 2020). The second is microenvironmental heterogeneity, in which litter mixtures create more heterogeneous physical and nutritional environments that alter decomposer communities and enhance decomposition (e.g., Hättenschwiler et al. 2005; Hättenschwiler and Gasser 2005; Liu et al. 2020).

However, both the direction and magnitude of litter-mixture effects vary widely depending on litter species composition, species richness, climatic conditions, litterbag mesh size (and thus the degree of soil-fauna involvement), and the duration of decomposition (e.g., Liu et al. 2020). Consequently, it remains important to evaluate litter-mixture effects for dominant plant species within individual forest ecosystems. Moreover, understanding the mechanisms underlying litter-mixture effects requires more than measuring decomposition of litter mixtures as a whole. By separating each litter species from mixed-species litterbags and comparing its decomposition with that in single-species litterbags, changes in the decomposition trajectory of individual litter species can be evaluated.

In cool-temperate deciduous forests of Japan, particularly in snowy regions, the forest floor is commonly dominated by dense understory dwarf bamboo (Sasa) species. As evergreen understory plants, Sasa species form a distinct understory layer beneath deciduous tree canopies and are important primary producers during periods when the forest floor receives sufficient light, particularly between autumn leaf fall and snow cover and between snowmelt and canopy leaf-out. Sasa leaves often have a lifespan of several years (for example, approximately two years in *Sasa senanensis*), with older leaves being shed mainly in autumn (Watanabe et al. 2013). In addition, Sasa litter is known to differ in both chemical characteristics and decomposition dynamics from deciduous tree litter. (e.g., Watanabe et al. 2013). In addition, Sasa litter constitutes a substantial proportion of annual litterfall, accounting for approximately 29–33% of total litter inputs in several forests (Tripathi et al. 2006; Watanabe et al. 2013).

One of the pioneering studies examining litter-mixture effects involving Sasa was conducted by Kaneko and Salamanca (1999). They reported that mixtures containing Sasa litter decomposed faster than expected from the component species alone, indicating synergistic, non-additive effects. They attributed this acceleration primarily to increased microhabitat complexity created by mixing morphologically different litter types, which promoted higher abundances of microarthropods and greater species richness of oribatid mites. However, studies investigating litter mixtures involving Sasa remain extremely limited. Similarly, Shi et al. (2015) reported accelerated decomposition in mixtures of Moso bamboo and evergreen broad-leaved litter in subtropical bamboo forests. However, because decomposition was evaluated only for the litter mixture as a whole, their study could not determine how species mixing affected the decomposition of each litter species or the associated changes in litter chemistry and microbial communities. Therefore, it remains unclear how mixing Sasa litter with deciduous tree litter influences not only the decomposition rate but also the decomposition trajectory of individual litter species.

Accordingly, this study aimed to clarify how litter mixtures of two dominant canopy tree species (oak and birch) and the dominant understory bamboo (Sasa) influence litter decomposition in cool-temperate deciduous forests in Japan. To this end, we quantified litter mass loss of each species in both single-species and mixed-species litterbags, together with changes in litter chemistry and microbial communities during decomposition. We then compared the decomposition trajectories of each litter species between single-species and mixed-species conditions to determine whether species mixing alters not only the rate but also the trajectory of litter decomposition.

## Materials and Methods

### Study site

This study was conducted from December 2014 to November 2016 in in a cool-temperate forest located on the northwest slope of Mount Norikura in Gifu Prefecture, Japan (Takayama Forest, 36°08′ N, 137°25′ E, 1420 m a.s.l.) near the Takayama Field Station of Gifu University, Japan. The mean annual air temperature and precipitation from 2000 to 2018, as recorded at the Takayama Field Station (ca. 300 m from the Takayama Forest), were 7.2°C and 2146 mm, respectively. This area is usually covered by snow from late December to early April.

Monthly air temperatures during the experimental period ranged from −4.8°C in January 2015 to 19.3°C in August 2016. Yearly precipitation was approximately 2270 and 2030 mm in 2015 and 2016, respectively, whereas winter precipitation from December 2014 to March 2015 and from December 2015 to March 2016 (falling mainly as snow) was approximately 800 and 470 mm, respectively. The maximum snow depth during each period was 205 and 73 cm, respectively.

This site is a secondary broad-leaved deciduous forest (approximately 50–60 years old) and has a permanent plot of 1 ha (100 m × 100 m). The forest was dominated by oak (*Quercus crispula*, 26.9% in total basal area) and birches (*Betula ermanii* (24.6%) and *B. platyphylla* var. *japonica* (14.6%)) (Ohtsuka et al. 2005). Although the forest floor is generally covered by dense (ca. 40 stems m^−2^) dwarf bamboo grass (*Sasa senanensis*) at 1–1.5 m height, some small patches without Sasa cover occurred around the permanent plot. The soil at the study site was classified as an andisol (Japanese volcanic ash soils) (Chen et al. 2017).

### Litter bag experiment

A two-year litter bag experiment (from December 2014 to November 2016) was conducted to examine how species mixing alters the decomposition rate and trajectory of each species litter in the area (ca. 30 m × 30 m) without Sasa cover near the permanent plot in the Takayama Forest. In November 2014, fresh leaf litter of oak (O, *Q. crispula*), birch (B, *B. ermanii*) and understory Sasa (S, *S. senanensis*) was collected around the permanent plots and air-dried. We designed 7 types of litter bags (single species (type O, B, and S), two-species mixture (type OB, OS, and BS), and three-species mixture (type OBS)) and prepared 42 bags for each bag type (294 bags in total). Numbered polyethylene mesh bag (bag size: 18 cm × 25 cm, mesh opening: ca. 2 mm) were filled with approximately 3.0 g of air-dried litter. Multi-species mixture bags also contained 3.0 g of air-dried litter samples in total with equal weight for each species. We estimated the oven-dry weight of litter in the bag on the ratio of the weight of the subsample before and after oven drying at 70°C. Extra subsamples of each species was stored for analyses of the initial chemical and microbial properties. On 7 December 2014, all litter bags were placed on the forest floor and were covered roughly by a polyethylene net (2.5 cm mesh) to prevent them from moving away by the snowmelt or from being disturbed by wild animals.

Six litter bags per each bag type were collected (42 bags in total) after 53, 95, 142, 275, 366, 436, and 701 days. After removing exogeneous materials such as soil particles or plant roots by hand using tweezers, litter samples were separated into each species as far as possible. The samples were freeze-dried, weighed, and stored at −80°C until further analysis.

Litter mass remaining (% of initial mass) at time *t* (*MR*_*t*_,) was calculated by dividing the dry weight of the remaining litter at time *t* (*W*_*t*_) by the initial dry weight of the litter (*W*_*i*_).

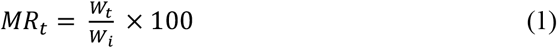

The sequential decrease in litter mass remaining during the ca. two years experiment was fitted by an exponential model (e.g., Olson 1963; Harmon et al. 2009):

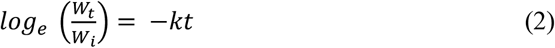

where *t* is time since litterbag placement (year), *k* is decomposition rate constant.

Accumulated litter mass loss at time *t* (*ML*_*t*_, % of initial mass) was expresses as follows:

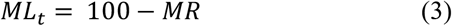

The effect of species mixing on litter decomposition was quantified as the natural log-transformed response ratio (ln R) by comparing the observed total mass loss of mixed-species litter bags (observed value, ML_obs_, %) with the expected mass loss (*ML*_*exp*_, %) calculated from the corresponding single-species-species litter bags. Positive values of ln R indicate synergistic effects, negative values indicate antagonistic effects, and zero indicates additive effects.

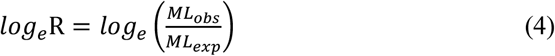

The *ML*_*exp*_ was calculated from the results in single-species litter bags as follows;

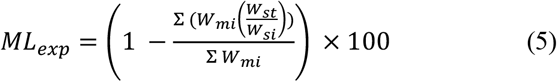

where *W*_*si*_ and *W*_*st*_ are the initial and remaining dry weights, respectively, of each litter species in the corresponding single-species litter bags, and *W*_*mi*_ is the initial dry weight of each litter species in the mixed-species litter bag.

### Chemical analyses

Dried litter samples were ground into powder using a mechanical mill (IFM-800DG; Iwatani Corporation, Tokyo, Japan) followed by grinding with a mortar and pestle to pass through a 500μm mesh sieve and used for subsequent chemical and microbial analyses. The total C and N contents of the samples were determined with a CN analyzer (Sumigraph NC-22; Sumika Chemical Analysis Co., Tokyo, Japan).

Acid-unhydrolyzable residue (also known as the acid-insoluble residue or ‘‘Klason lignin’’ fraction) of initial litter sample and ca. one-year (366 days) and two-year (701 days) litter bag samples were determined by the gravimetric method with alcohol-benzene extraction and sulfuric acid digestion according to the method of King and Heath (1967) as described in Osono et al. (2014) with some modification. Powdered litter sample was extracted with ethanol-benzene (1:2, v/v) using Soxhlet extractors for 6–8 hours. Then, 30 mg of the air-dried residue was treated with 72% sulfuric acid at 30°C for 2.5 hours. Next, the mixture was diluted to make a 2.5 % sulfuric acid solution and autoclaved at 120°C for 60 min. After cooling, the residue was filtered with Whatman grass fiber filter (GF/C), dried at 105°C, and weighed as acid-insoluble residue. In this study, this acid-insoluble residue (Klason lignin) was hereafter referred to as lignin.

Silicate concentration in the initial litter samples was determined by the gravimetric method with wet digestion as described in Watanabe et al. (2014) with some modification. The powdered 250 mg samples was digested with 60% nitric acid and 30% hydrogen peroxide and the residue in the digested solution was obtained by filtrating through filter paper (Advantec No. 6; Tokyo, Japan), followed by burning at 550°C for 2 h and 1000°C for 5 min, and weighing as silicate (SiO_2_).

Element or component remaining at time *t* (*ER*_*t*_, % of initial weight) for total C, total N, and lignin was calculated by dividing the element or component weight of the remaining litter (*EW*_*t*_) by the initial weight (*EW*_*i*_):

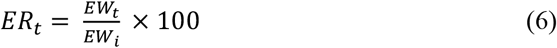

### Microbial analyses

Microbial biomass and community structure in the decomposing litter were examined by phospholipid fatty acid (PLFA) analysis. PLFA is the main component of living cell membranes, and its composition is known to differed among taxonomically different microbes; therefore, the amount and composition of PLFA have been widely used as an index of the microbial biomass and as a fingerprint of the taxonomical microbial community structure, respectively (e.g., Frostegård et al. 1993; White et al. 1979; Yoshitake et al. 2020).

We followed the basic procedure for PLFA analysis as described in Yoshitake et al. (2020). Briefly, total lipids were extracted in a chloroform–methanol–citrate buffer from a freeze-dried 100–500 mg powdered litter sample; the lipids were separated on a silicic acid column, and only the phospholipid fraction was obtained. The phospholipids were esterified and the resultant fatty acid methyl esters were separated using a gas chromatograph (GC 2014; Shimadzu Corp., Kyoto, Japan). Peak areas were quantified against methyl nonadecanoate fatty acid (19:0) as an internal standard. The total PLFA content (μmol g^−1^) was used as an index of total microbial biomass (Frostegård et al. 1991). The sums of nine fatty acids (i15:0, a15:0, 15:0, i16:0, 17:0, i17:0, cy17:0, 18:1ω7 and cy19:0) were used as bacterial PLFA (μmol g^−1^) (e.g., Federle 1986; Frostegård et al. 1993), whereas 18:2ω6 was used as fungal PLFA (μmol g^−1^) (e.g., Federle 1986; Frostegård et al. 1993). The fungal-to-bacterial PLFA ratio (F/B ratio) was then calculated.

### Statistics

One-way analysis of variance (one-way ANOVA) and Tukey’s test were applied to test for the significant differences in initial chemical properties among species.

Decomposition constants (*k*) were estimated as the slopes of origin-constrained regressions of log-transformed litter mass remaining against time (Eq. 3). Differences in *k* among groups were assessed by comparing group-specific slopes within a linear model, followed by Tukey-adjusted pairwise comparisons.

To characterize and compare decomposition trajectories, relationships between mass loss (*ML*_*t*_, % of initial mass) and element/component remaining (*ER*_*t*_) were examined using regression analyses. For each group, both linear and quadratic regression models were fitted and model performance was evaluated using the adjusted *R*^2^, the statistical significance of regression coefficients (*p*) and Akaike Information Criterion (AIC). The most appropriate model form (linear or quadratic) was selected based on these criteria.

Differences in regression relationships were evaluated by comparing nested models with and without interaction terms using ANOVA. When a significant interaction was detected, pairwise comparisons of regression slopes were performed using estimated marginal trends with Tukey’s adjustment for multiple comparisons.

Species-mixture effects were evaluated using the log-transformed response ratio (ln R, eqn (4)). A value of ln R = 0 indicates no species-mixture effect. For each bag type, we tested whether mean ln R differed significantly from zero using a one-sample t-test.

Differences in PLFA contents and F/B ratio of individual litter species in different types of litter bags were tested by one-way analysis of variance (one-way ANOVA) and Tukey’s test.

All statistical analyses were performed in R (version 4.5.1; R Development Core Team 2025), and significance was evaluated at *p* < 0.05.

## Results

### Initial chemical properties of leaf litter

Initial C and N contents and C/N ratio differed significantly among three plant species (*p* < 0.05; Table 1). Oak leaf litter was characterized by the lowest N content and the highest C/N ratio, whereas birch showed the highest N content and the lowest C/N ratio. Sasa leaf litter showed intermediate N content and C/N ratio. Significant differences in lignin and silicate contents were observed between the deciduous tree species (oak and birch) and Sasa (Table 1). Sasa litter was characterized by significantly lower lignin and higher silicate contents.

**Table 1.** Chemical properties of leaf litter used for the litterbag experiment. Values are presented as means (SD; *n* = 6 for total C, N, and C/N ratio; *n* = 3 for lignin, lignin/N ratio, and Si). Values with different letters differ significantly (Tukey’s test, *P* < 0.05).

|  |  | Oak |  |  | Birch |  |  | Sasa |  |  |
| --- | --- | --- | --- | --- | --- | --- | --- | --- | --- | --- |
| Total C | (%) | 49.1 | (0.326) | b | 50.3 | (0.381) | c | 41.5 | (0.847) | a |
| Total N | (%) | 0.712 | (0.0247) | a | 1.08 | (0.0302) | c | 0.782 | (0.0689) | b |
| C/N ratio |  | 69.1 | (2.30) | c | 46.6 | (1.25) | a | 53.3 | (4.21) | b |
| Lignin | (%) | 44.8 | (4.21) | b | 43.3 | (2.08) | b | 25.9 | (3.73) | a |
| Lignin/N ratio |  | 63.3 | (6.03) | b | 39.9 | (2.43) | a | 30.9 | (4.57) | a |
| Si | (%) | 0.0500 | (0.00874) | a | 0.0670 | (0.0383) | a | 4.91 | (2.66) | b |

### Temporal changes in litter decomposition in single-species litterbags

Litter mass remaining in the single-species litterbags (types O, B, and S) gradually decreased over time (Fig. 1a). After two years, mass remaining of Sasa litter was significantly higher (about 60%) than those of oak (48%) and birch (44%). In addition, the decomposition rate constant (*k*) of Sasa litter (0.240) was significantly lower than those of deciduous tree species (0.430 for oak and 0.407 for birch).

**Fig. 1.**
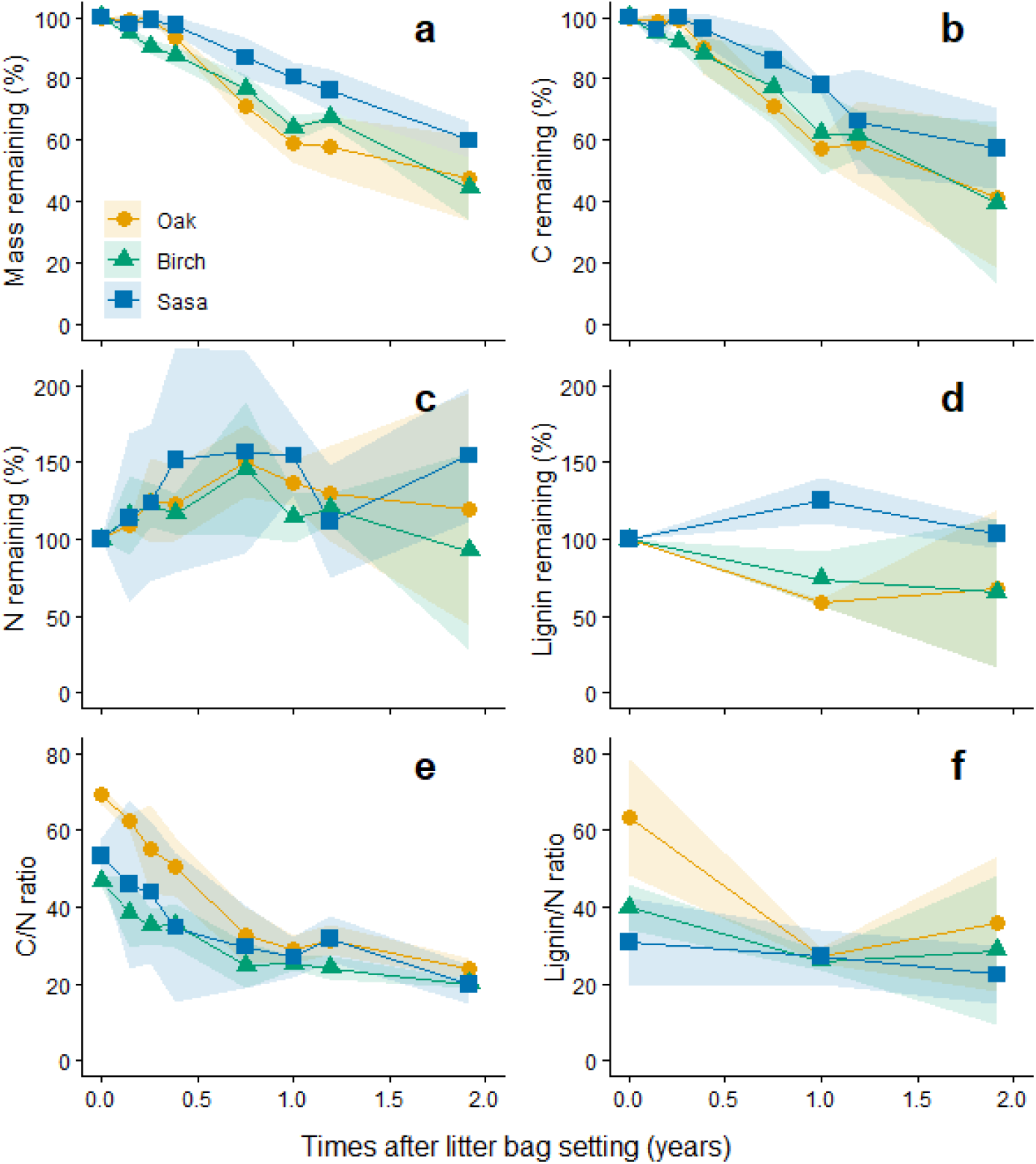
Changes over time in (a) mass remaining, (b) carbon remaining, (c) nitrogen remaining, (d) lignin remaining, (e) C/N ratio, and (f) lignin/N ratio of litter from three plant species (O, oak; B, birch; S, Sasa) in single-species litterbags over the two-year decomposition experiment. Points and shaded areas represent means and 95% confidence intervals, respectively (*n* = 6 litterbags at each sampling time).

Change in C remaining over time (% of initial weight) was similar to that in mass remaining (Fig. 1b). Unlike C remaining, N remaining increased to about 140–160% of the initial value at the end of the first year and then turned to decline in the second year for all species (Fig. 1c). However, N remaining in Sasa litter increased again during the second year. As a result, litter N remaining of deciduous tree species at the end of second year were about 90–120%, whereas it was still over 150% for Sasa litter (Fig. 1c). Lignin remaining for deciduous tree species decreased to about 60–70% of initial but the remaining in Sasa litter did not decrease (Fig. 1d). Litter C/N ratio, which originally differed significantly among species, continued to decrease from the beginning of experiment and reached similar values (about 25–30) at the end of the first year for all species (Fig. 1e). Thereafter the C/N ratio was relatively stable or slightly decreased in the second year. Litter lignin/N ratio also decreased rapidly in the first year and differences among species became smaller (Fig. 1f).

### Decomposition trajectories in single-species litterbag

To compare decomposition trajectories among plant species independently of differences in decomposition rate, changes in element/component remaining were examined against litter mass loss rather than time after litterbag placement (Fig. 2). Based on the model selection results (Table A1), linear regression models were selected for all species for C remaining (Fig. 2a) but the relationships did not differ significantly among the three species (*p* = 0.640).

**Fig. 2.**
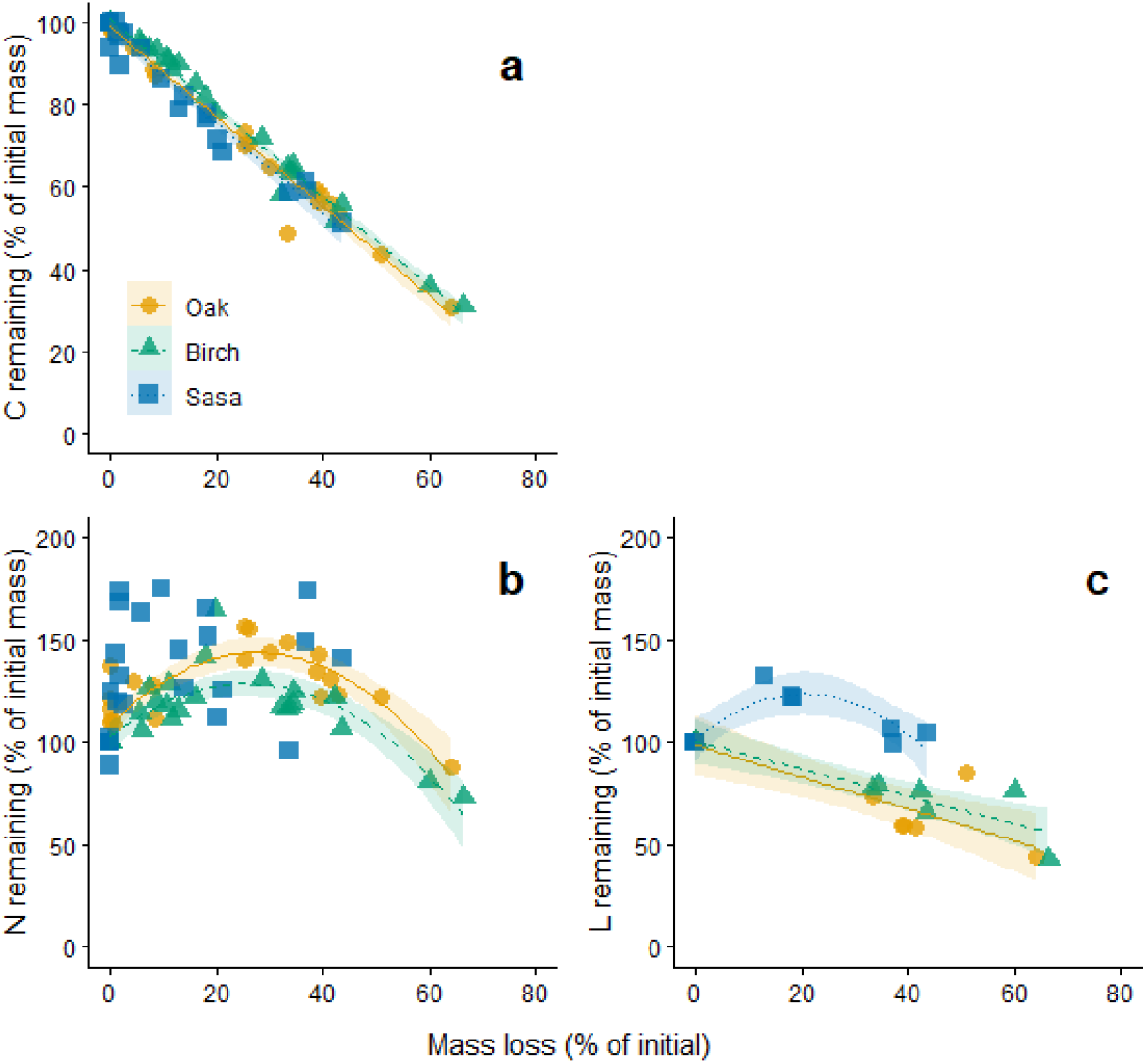
Decomposition trajectories showing relationships between litter mass loss and (a) carbon remaining, (b) nitrogen remaining, and (c) lignin remaining of litter from three plant species (O, oak; B, birch; S, Sasa) in single-species litterbags. Each point represents an individual litterbag, and shaded areas represent the 95% confidence intervals of the fitted regression lines.

For N remaining, quadratic regression models were selected for oak and birch, whereas no significant regression model was identified for Sasa (Fig. 2b; Table A1). Fitted relationships did not differ significantly between oak and birch (*p* = 0.376).

For lignin remaining, linear regression models were selected for oak and birch, whereas a quadratic regression model was selected for Sasa (Fig. 2c; Table A1). The linear regression relationships did not differ significantly between oak and birch (*p* = 0.610).

### Microbial community in single-species litterbag

Total microbial biomass (total PLFA content) increased rapidly after litterbag placement in all species (Fig. 3a). A marked increase in fungal biomass was also observed mainly in the first half of the first year and thereafter it decreased to approximately one-third of the maximum value at the end of the second year (Fig. 3b). Bacterial biomass increased more slowly than fungal biomass and continued to increase during the second year except for Sasa where it decreased during the second year (Fig. 3c). Although the fungal-to-bacterial PLFA ratio (F/B ratio) was initially much higher in the deciduous tree species than in Sasa, it declined markedly during the first year and approached the value observed in Sasa litter (Fig. 3d).

**Fig. 3.**
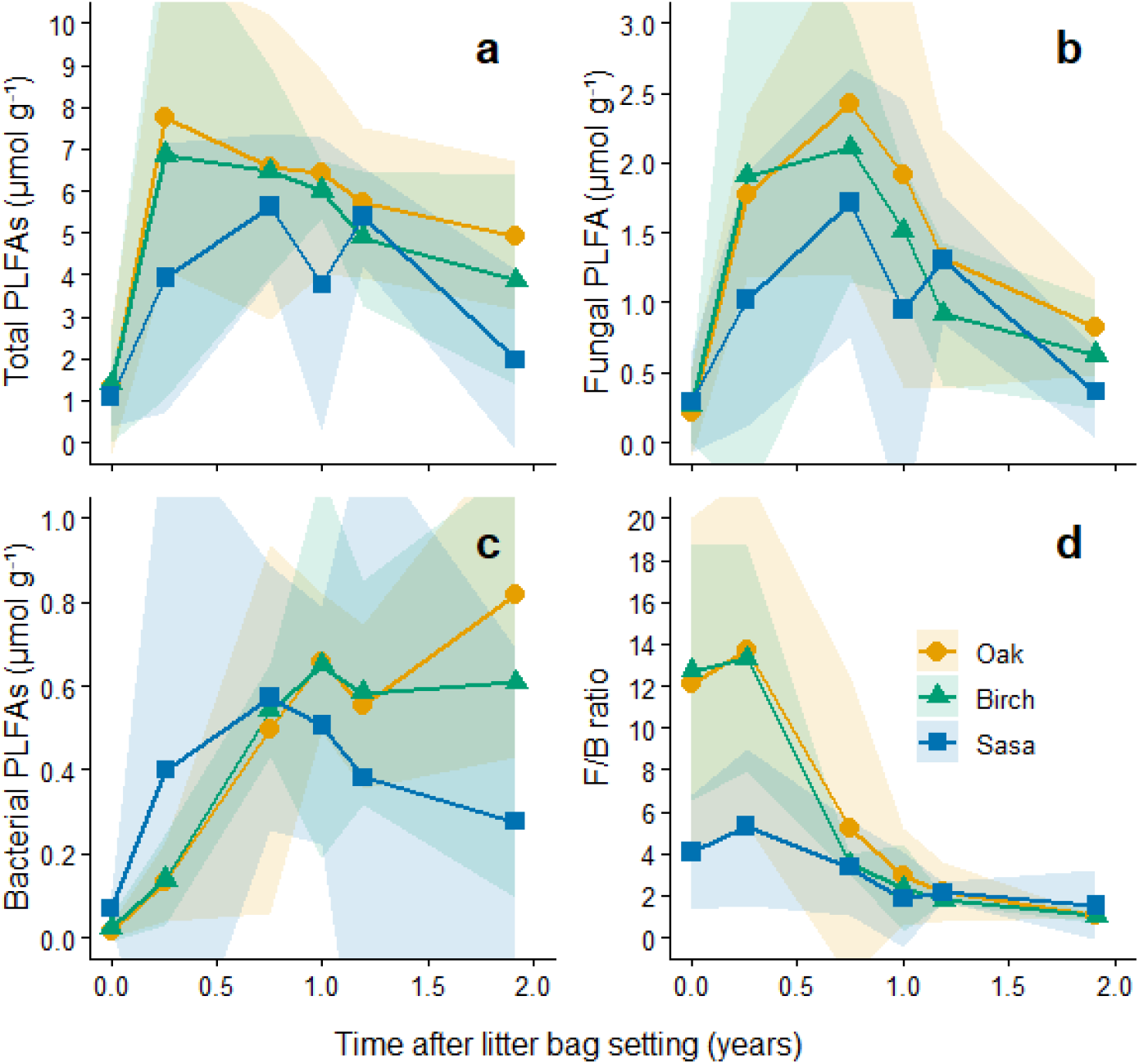
Changes over time in (a) total, (b) fungal, and (c) bacterial PLFA contents, and (d) the fungal-to-bacterial PLFA ratio (F/B ratio), of litter from three plant species (O, oak; B, birch; S, Sasa) in single-species litterbags over the two-year decomposition experiment. Points and shaded areas represent means and 95% confidence intervals, respectively (*n* = 3 litterbags at each sampling time).

### Effect of species mixing on mass loss

The log-transformed response ratio (ln R) did not differ significantly from zero for any mixed-species litterbag type (one-sample t-test, Fig. 4). However, ln R for OB tended to be greater than zero (*p* = 0.260), indicating that the observed mass loss exceeded the expected value calculated from the corresponding single-species litterbags (i.e., a synergistic effect). In contrast, the log R for other bag types (OS, BS, OBS) tended to be less than zero, indicating a tendency toward antagonistic effects (*p* = 0.162, 0.0583, and 0.103 for OS, BS, and OBS, respectively).

**Fig. 4.**
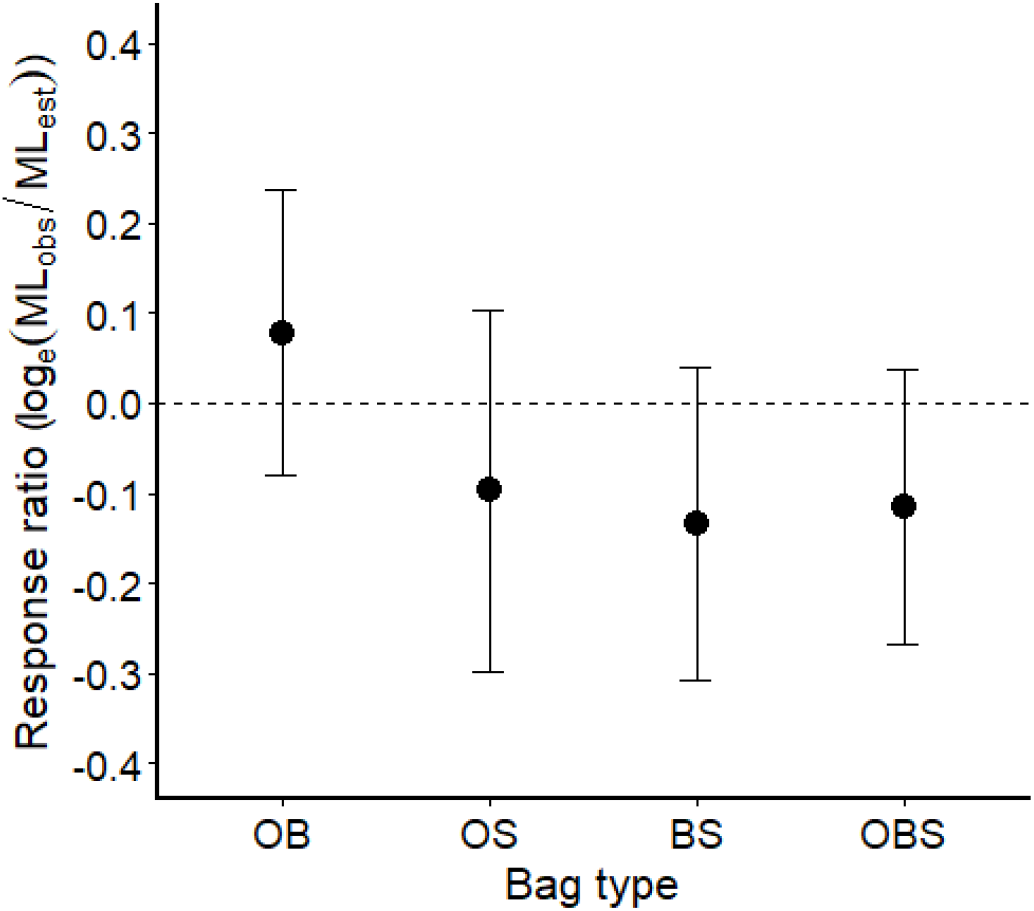
Natural log-transformed response ratio of observed to expected litter mass loss for mixed-species litterbags (ln R). Expected mass loss was calculated based on the mass remaining of each litter species in single-species litterbags (Eq. 5). Points and error bars represent means and 95% confidence intervals, respectively (*n* = 6).

### Effect of species mixing on decomposition trajectories

Based on the model selection results (Table A2), linear regression models were selected for C remaining in all species and litter-mixture treatments (Fig. 5a−c). Only for Sasa, regression slopes differed significantly among litter-mixture treatments, with the slope for the three-species mixture (OBS) being significantly steeper than those for the other three litter-bag types (Tukey’s test, *p* < 0.01, Fig. 5c).

**Fig. 5.**
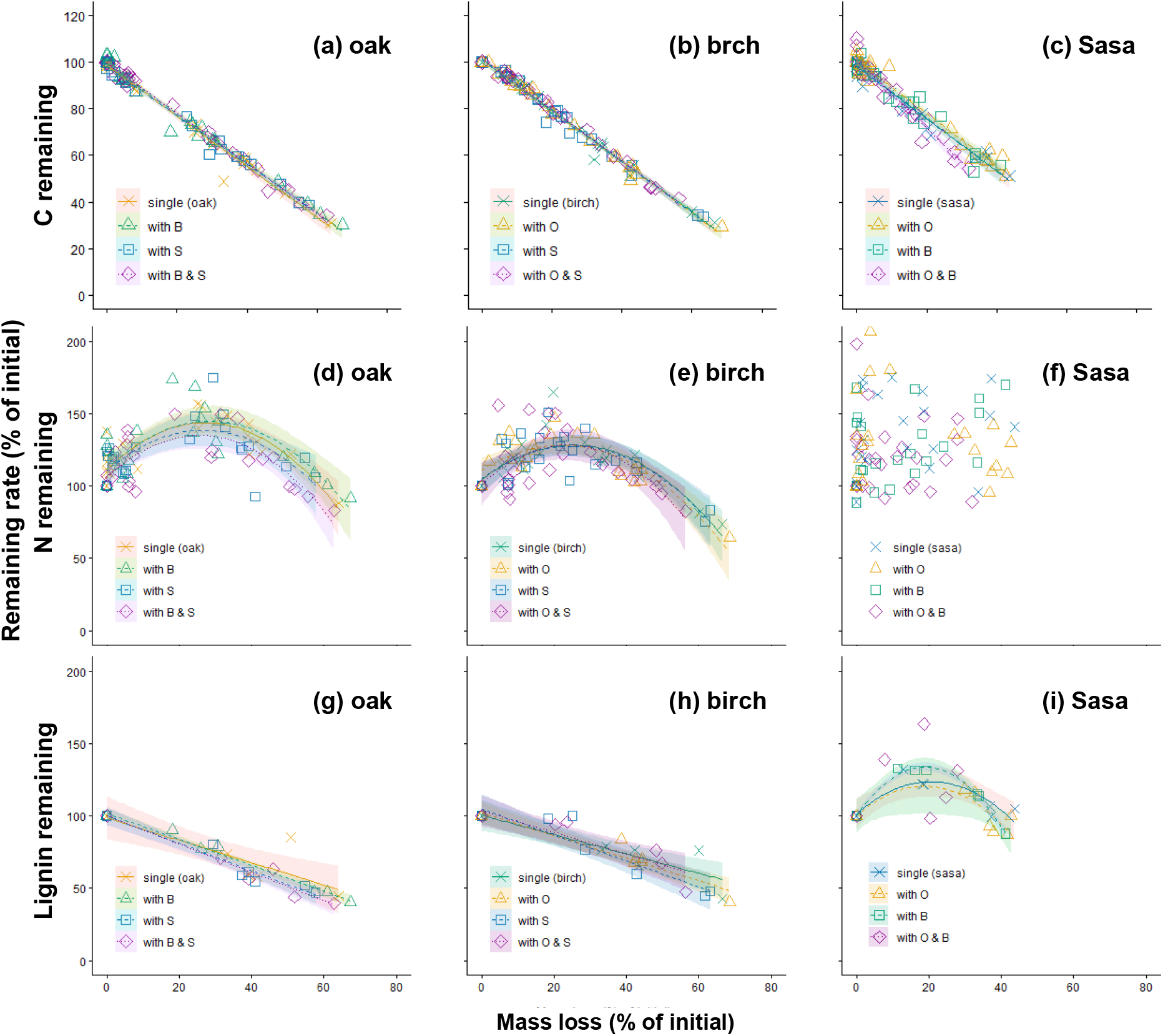
Decomposition trajectories showing the relationships between litter mass loss and carbon remaining (a–c), nitrogen remaining (d–f), and lignin remaining (g–i) for oak (a, d, g), birch (b, e, h), and Sasa (c, f, i) in single-species and mixed-species litterbags. Each point represents an individual litterbag, and shaded areas represent the 95% confidence intervals of the fitted regression lines.

For N remaining, quadratic regression models were selected for all litter-mixture treatments in oak and birch, whereas no significant regression model was obtained for Sasa (Table A2; Fig. 5d−f). In both oak and birch, the regression relationships did not differ significantly among litter-mixture treatments (Fig. 5d,e).

For lignin remaining, linear regression models were selected for all litter-mixture treatments in oak and birch (Fig. 5g,h), whereas quadratic regression models were selected for Sasa except for the three-species mixture treatment (OBS), for which no significant regression models were obtained (Fig. 5i). Within each species, the regression relationships did not differ significantly among litter-mixture treatments.

Despite these largely conserved decomposition trajectories, decomposition rate constants (*k*) showed treatment-dependent trends (Table 2). Although not always statistically significant, the decomposition constant (*k*) of deciduous tree species (oak and birch) tended to be higher when litter was mixed with each other (OB treatment) but lower when mixed with Sasa litter (OS, BS, and OBS treatments) than in single-species litter bags. In contrast, the *k* value of Sasa tended to decrease in treatments containing birch litter (BS and OBS).

**Table 2.** Decomposition constants (*k*) for leaf litter of three species obtained from single-species and species-mixed litterbags. Values with different letters differ significantly (Tukey-adjusted pairwise comparisons). ns, not significant.

| | $k$ (yr<sup>-1</sup>) | |
| --- | --- | --- |
| Oak (O) |  |  |
| Single | 0.43 | ns |
| with B | 0.44 | ns |
| with S | 0.40 | ns |
| with B & S | 0.42 | ns |
| Birch (B) |  |  |
| Single | 0.41 | ab |
| with O | 0.47 | a |
| with S | 0.35 | b |
| with O & S | 0.36 | b |
| Sasa (S) |  |  |
| Single | 0.24 | a |
| with O | 0.24 | a |
| with B | 0.21 | ab |
| with O & B | 0.18 | b |

### Effect of species mixing on microbial community

For deciduous tree species (oak and birch), total microbial biomass (total PLFA content) tended to be lower in mixed-species litterbags at the end of the first year (Fig. 6a, b), but no significant differences were detected among treatments at the end of the second year. In contrast, microbial biomass in Sasa litter tended to be higher in mixed-species litterbags at the end of the second year (Fig. 6c).

**Fig. 6.**
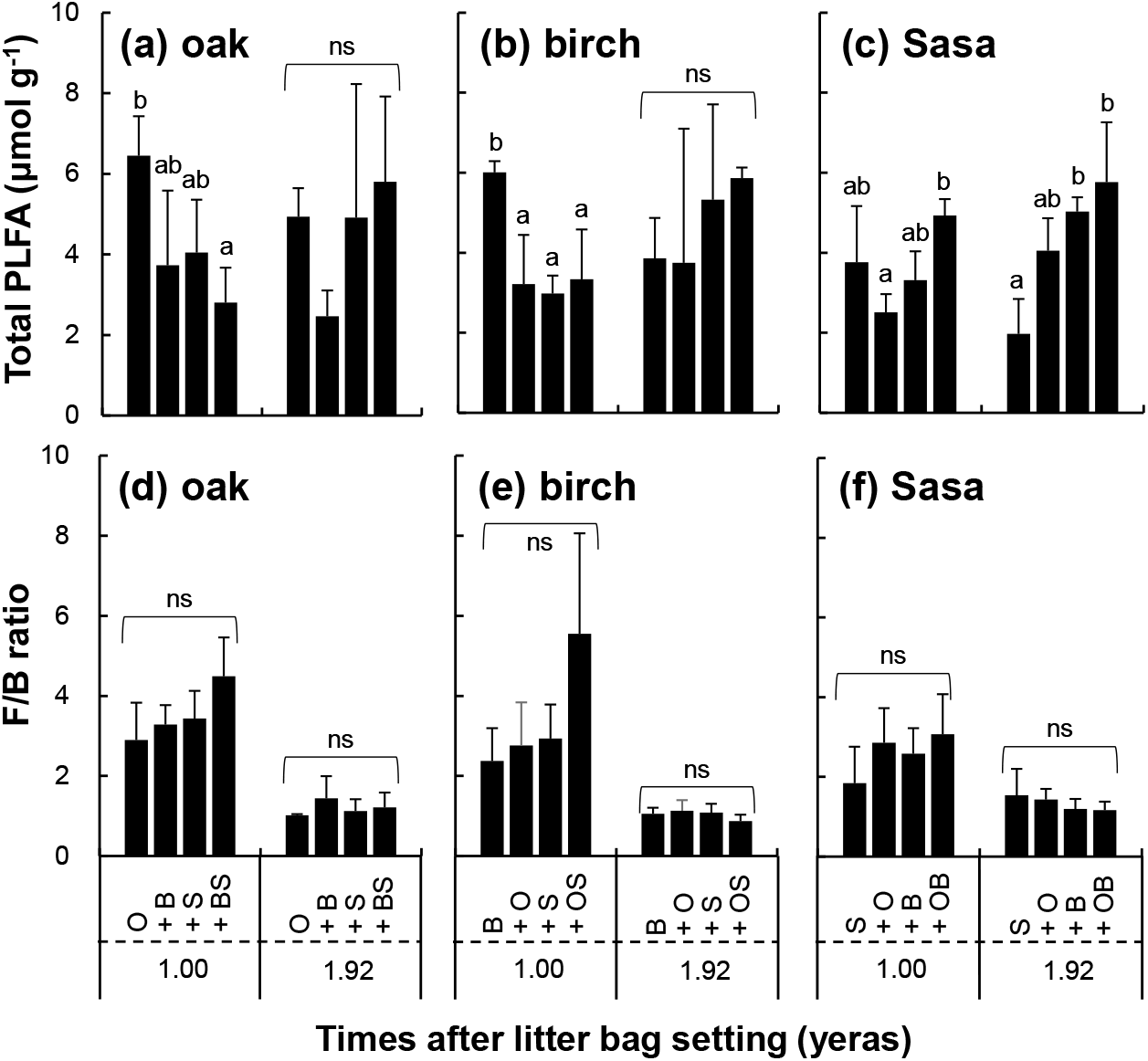
(a–c) Total PLFA content and (d–f) fungal-to-bacterial PLFA ratio (F/B ratio) of decomposing litter from oak (a, d), birch (b, e), and Sasa (c, f) in single-species and mixed-species litterbags after one and two years of decomposition. Values are means and error bars represent standard deviation, respectively (*n* = 3). Different letters indicate significant differences among litterbag treatments within each sampling year (Tukey’s test, *p* < 0.05). ns, not significant.

The F/B ratio showed no significant differences among litter-mixture treatments for any species, although oak and birch tended to exhibit slightly higher values in the OBS treatment at the end of the first year (Fig. 6d−f).

## Discussion

Litter decomposition of deciduous broad-leaved tree species (oak and birch) was faster than that of understory Sasa in single-species litter bag (Fig. 1a). Observed *k* value for oak (0.430), birch (0.407), and Sasa (0.240) in this study (Table 2) were all within the ranges previously reported for the same species or genius (Salamanca et al. 1998; Kaneko and Salamanca 1999; Osono and Takeda 2005; Miyamoto and Hiura 2008; Osono et al. 2014). Significantly slower decomposition of Sasa litter which had intermediate C/N ratio and lowest lignin/N ratio would be related to the high concentration of silicate (c.f. Table 1) (Kaneko and Salamanca 1999, Watanabe et al. 2013). For all three species, C content decreased with mass loss, whereas N content once increased due to microbial N immobilization, during which microbes grow (construct their body) while consuming organic C in litter and absorbing inorganic N in and out of litter (e.g., Melillo et al. 1982; Osono et al. 2014), and then decreased (Fig. 2a, b). As a result of changes in C and N contents, C/N ratio and lignin/N ratio rapidly decreased in the first year (Fig. 1e, f).

Changes in total microbial biomass showed broadly similar temporal patterns among the three litter species. Total microbial biomass increased rapidly during the early stage of decomposition in all litter types (Fig. 3a), largely owing to the rapid increase in fungal biomass (Fig. 3b). This pattern is consistent with previous studies showing that fungi play a dominant role during the initial stage of leaf litter decomposition

(e.g., Osono and Takeda 2001; Lodato et al. 2021). In contrast, bacterial biomass increased more gradually and continued to increase during the second year in the deciduous tree litter (Fig. 3c), suggesting a greater contribution of bacteria during the later stage of decomposition.

Interestingly, the fungal-to-bacterial PLFA ratio (F/B ratio) during the early stage of decomposition was approximately three times lower in Sasa litter than in the two deciduous tree species (Fig. 3d), indicating a lower relative abundance of fungi. Given that fungi play a crucial role in the decomposition of fresh leaf litter, this lower fungal dominance may have contributed to the slower decomposition of Sasa litter.

Together with the distinct patterns of N and lignin dynamics, these microbial characteristics are consistent with the distinct decomposition trajectory observed for Sasa litter.

Although these temporal changes generally agreed with existing knowledge, examining changes as a function of litter mass loss rather than elapsed time revealed an additional pattern that was not apparent from time-course analyses alone. Specifically, decomposition trajectories were largely conserved between the two deciduous tree species (oak and birch) despite differences in their initial litter chemistry. There were no significant differences in the relationships between mass loss and C, N, or lignin remaining for these two species (Fig. 2). This suggests that, once decomposition begins, deciduous broad-leaved litter follows a common decomposition trajectory regardless of species identity.

In contrast, Sasa exhibited a distinct decomposition trajectory compared with the two deciduous tree species, showing no significant relationship between N remaining and mass loss (Fig. 2b) and a markedly different pattern of lignin dynamics (Fig. 2c). These results indicate that Sasa litter is not simply decomposed more slowly than deciduous tree litter; rather, it follows a fundamentally different decomposition trajectory.

Species mixing has often been discussed in terms of additive versus non-additive effects on litter decomposition (e.g., Liu et al. 2020; Bai et al. 2026). In the present study, the response ratio (ln R) did not differ significantly from zero for any litter mixture (Fig. 4), indicating that overall litter decomposition was largely additive. Nevertheless, mixtures of the two deciduous tree species (OB) tended to accelerate decomposition (ln R > 0), whereas mixtures including Sasa tended to reduce decomposition rates (ln R < 0) (Fig. 4). To clarify how these mixture effects arose, we examined decomposition of each litter species separately. The decomposition rate constant (*k*) of oak and birch tended to increase when mixed with each other but decrease when mixed with Sasa (Table 2). Likewise, decomposition of Sasa tended to be slower in mixtures containing birch. Despite these changes in decomposition rate, decomposition trajectories remained remarkably similar among litter-mixture treatments within each species (Fig. 5). Thus, litter mixture altered how rapidly decomposition proceeded, but had little influence on the underlying decomposition trajectory of each litter species.

Changes in decomposition rate were partly accompanied by changes in microbial biomass. Actually, mixing with Sasa litter reduced microbial biomass in oak and birch litter (Fig. 6), which likely contributed to the slower decomposition rate. Because the F/B ratio did not change, the suppression appeared to be associated mainly with microbial abundance rather than changes in the fungal-to-bacterial balance. In contrast, decrease in decomposition rate (*k*) of Sasa litter in the litter bags including birch litter (BS and OBS, Table 2) cannot be fully explained by significantly increased microbial biomass and stable F/B ratio (Fig. 6). Several, not mutually exclusive, mechanisms may explain this discrepancy. Increased microbial biomass does not necessarily indicate higher microbial activity. In addition, similar F/B ratios do not necessarily indicate similar microbial community composition because microbial taxa within fungal and bacterial groups may differ among treatments. Future studies should therefore examine microbial enzyme activities and/or microbial community composition at a finer taxonomic resolution to better understand the mechanisms underlying litter-mixture effects.

Previous studies have shown that litter-mixture effects involving Sasa are not uniform. Kaneko and Salamanca (1999) reported accelerated decomposition in mixtures of Sasa (*Sasa veitchii*) with pine (*Pinus densiflora*) litter, whereas decomposition of oak (*Quercus serrata*) litter was little affected by Sasa. They suggested that mixing morphologically contrasting litter types increased habitat heterogeneity, thereby enhancing the abundance and richness of litter microarthropods and promoting decomposition. In the present study, however, all litter species were broad-leaved and therefore differed less in physical structure than the pine–Sasa combination examined by Kaneko and Salamanca (1999). Consequently, litter mixtures may have generated only limited structural heterogeneity, reducing the potential for habitat-mediated facilitation of decomposer communities. This interpretation is also consistent with our PLFA results, which provided little evidence that litter mixing consistently enhanced microbial biomass or altered the fungal-to-bacterial balance.

In contrast to the accelerated decomposition reported for pine–Sasa mixtures, mixtures including Sasa in the present study tended to reduce decomposition rates of deciduous tree litter. Together, these findings suggest that the effects of Sasa on litter decomposition depend not only on litter quality but also on the physical characteristics of the accompanying litter species, rather than representing a universal property of Sasa litter itself.

Although litter-mixture effects varied depending on the species combination, decomposition trajectories were largely conserved within each litter species (Fig. 5). This suggests that species mixing primarily influences the rate at which decomposition proceeds, whereas the sequence of chemical transformations during decomposition is an intrinsic property of each litter type.

## Conclusions

Cool-temperate forests in Japan are commonly characterized by a dense understory of Sasa, which contributes a substantial proportion of annual litter input. Owing to its distinctive litter chemistry, Sasa litter was associated with a distinct microbial community and decomposition trajectory compared with deciduous broad-leaved tree litter. Although mixing Sasa litter with deciduous tree litter tended to reduce decomposition rates, it did not alter the decomposition trajectories of the individual litter species. These findings suggest that Sasa influences litter decomposition primarily by modifying the pace, rather than the sequence, of decomposition processes.

From an ecological perspective, changes in the abundance or distribution of Sasa are therefore likely to influence litter decomposition and nutrient cycling not only through direct changes in litter input, but also indirectly through litter-mixture effects on decomposition rates. Such changes may occur following forest management practices in Satoyama landscapes (e.g., Tokumoto et al. 2026), decline of Sasa caused by herbivory (e.g., Fu et al. 2025; Yamada & Takatsuki 2015), or the synchronous flowering and subsequent die-off that periodically occur in many Sasa species (Abe et al. 2001; Yamada et al. 2023). Our results suggest that these vegetation changes may alter the timing of litter decomposition and nutrient turnover while leaving the decomposition trajectories of individual litter types largely unchanged.

## Acknowledgments

We thank Mr. K. Kurumado, Mr. Y. Miyamoto, Mr. Y. Hiomo, and K. Suzuki of the Takayama Experimental Field Station, Gifu University, Japan, for their great help and support at the station. We thank Dr. Nobuhide Fujitake and the members of the Soil Science Laboratory, Kobe University, for their technical assistance with the chemical analyses of litter samples.

## Author contributions

S. Y., C. A., and T. O. conceived the study and designed the methodology. C. A. conducted the investigation. C. A. and S. Y. contributed to data curation and visualization. S. Y. wrote the original draft. S. Y. and T. O. contributed to writing, review, and editing of the manuscript. All authors have read and approved the final version of the manuscript.

## Data availability

The data analyzed during the study are available from the corresponding author on request.

## Appendices

**Table A1.** Linear and quadratic regression models fitted to mass loss (ML, %) and element/component remaining (ER, %) data from single-species litter bags (Fig. 2).

| | Liner regression:<br>ER = $a$ ML + $b$ | | | Quadratic regression:<br>ER = $m(\text{ML}-n)^2 + o$ | | | Selected<br>regression<br>type | Equation |
| --- | --- | --- | --- | --- | --- | --- | --- | --- |
| | Adjusted<br>$R^2$ | $p$ | AIC$_l$ | Adjusted<br>$R^2$ | $p$<br>(quad) | AIC$_q$ | | |
| *Carbon remaining* |  |  |  |  |  |  |  |  |
| Oak | 0.981 | <0.001 | 141 | 0.982 | 0.136 | 140 | Linear | -1.09 [ML] + 99.0 |
| Birch | 0.988 | <0.001 | 123 | 0.988 | 0.723 | 125 | Linear | -1.08 [ML] + 101 |
| Sasa | 0.965 | <0.001 | 139 | 0.973 | 0.00590 | 132 | Linear | -1.13 [ML] + 98.4 |
| *Nitrogen remaining* |  |  |  |  |  |  |  |  |
| Oak | 0.0880 | 0.0726 | 237 | 0.697 | <0.001 | 209 | Quadratid | -0.0456 ([ML] - 27.7)<sup>2</sup> + 144 |
| Birch | -0.0130 | 0.423 | 237 | 0.656 | <0.001 | 208 | Quadratid | -0.0386 ([ML] - 25.8)<sup>2</sup> + 129 |
| Sasa | 0.0720 | 0.0953 | 259 | 0.107 | 0.170 | 259 | None | None |
| *Lignin remaining* |  |  |  |  |  |  |  |  |
| Oak | 0.713 | 0.00257 | 73.2 | 0.678 | 0.647 | 74.9 | Linear | -0.773 [ML] + 98.6 |
| Birch | 0.804 | <0.001 | 67.5 | 0.778 | 0.701 | 69.2 | Linear | -0.668 [ML] + 100 |
| Sasa | -0.142 | 0.961 | 75.7 | 0.639 | 0.00698 | 66.0 | Quadratic | -0.0524 ([ML] - 20.6)<sup>2</sup> + 124 |

**Table A2.** Linear and quadratic regression models fitted to mass loss (ML, %) and element/component remaining (ER, %) data from mixed-species litter bags (Fig. 5).

| | Liner regression:<br>ER = $a$ ML + $b$ | | | Quadratic regression:<br>ER = $m(\text{ML}-n)^2 + o$ | | | Selected<br>regression<br>type | Equation |
| --- | --- | --- | --- | --- | --- | --- | --- | --- |
| | Adjusted<br>$R^2$ | $p$ | AIC$_l$ | Adjusted<br>$R^2$ | $p$<br>(quad) | AIC$_q$ | | |
| Oak (O) |  |  |  |  |  |  |  |  |
| Carbon remaining |  |  |  |  |  |  |  |  |
| Single | <0.001 | 0.979 | 128 | 0.188 | 0.980 | 128 | Linear | -1.09 [ML] + 98.8 |
| with B | <0.001 | 0.982 | 125 | 0.0280 | 0.985 | 122 | Linear | -1.06 [ML] + 99.2 |
| with S | <0.001 | 0.991 | 106 | 0.284 | 0.991 | 106 | Linear | -1.05 [ML] + 98.3 |
| with B & S | <0.001 | 0.993 | 113 | 0.735 | 0.993 | 115 | Linear | -1.08 [ML] + 99.7 |
| Nitrogen remaining |  |  |  |  |  |  |  |  |
| Single | 0.227 | 0.0230 | 212 | <0.001 | 0.675 | 186 | Quadratic | -0.0437 ([ML] - 27.2)<sup>2</sup> + 144 |
| with B | 0.810 | -0.0430 | 221 | <0.001 | 0.538 | 203 | Quadratic | -0.0405 ([ML] - 28.4)<sup>2</sup> + 145 |
| with S | 0.351 | -0.00400 | 212 | 0.00129 | 0.365 | 202 | Quadratic | -0.0395 ([ML] - 26.7)<sup>2</sup> + 138 |
| with B & S | 0.558 | -0.0260 | 237 | <0.001 | 0.510 | 218 | Quadratic | -0.0442 ([ML] - 25.6)<sup>2</sup> + 135 |
| Lignin remaining |  |  |  |  |  |  |  |  |
| Single | 0.00257 | 0.713 | 73.2 | 0.647 | 0.678 | 74.9 | Linear | -0.773 [ML] + 98.6 |
| with B | <0.001 | 0.985 | 49.4 | 0.0868 | 0.989 | 46.6 | Linear | -0.897 [ML] + 102 |
| with S | <0.001 | 0.956 | 57.3 | 0.645 | 0.950 | 58.9 | Linear | -0.947 [ML] + 99.9 |
| with B & S | <0.001 | 0.967 | 50.9 | 0.992 | 0.960 | 52.9 | Linear | -0.977 [ML] + 100 |
| Birch (B) |  |  |  |  |  |  |  |  |
| Carbon remaining |  |  |  |  |  |  |  |  |
| Single | <0.001 | 0.987 | 112 | 0.489 | 0.987 | 114 | Linear | -1.09 [ML] + 101 |
| with O | <0.001 | 0.992 | 101 | 0.286 | 0.992 | 102 | Linear | -1.09 [ML] + 101 |
| with S | <0.001 | 0.987 | 110 | 0.419 | 0.987 | 112 | Linear | -1.09 [ML] + 101 |
| with O & S | <0.001 | 0.994 | 102 | 0.054 | 0.994 | 99.3 | Linear | -1.10 [ML] + 102 |
| Nitrogen remaining |  |  |  |  |  |  |  |  |
| Single | 0.153 | 0.0490 | 210 | <0.001 | 0.628 | 189 | Quadratic | -0.0373 ([ML] - 25.3)<sup>2</sup> + 128 |
| with O | 0.0404 | 0.140 | 206 | <0.001 | 0.661 | 185 | Quadratic | -0.0349 ([ML] - 22.3)<sup>2</sup> + 128 |
| with S | 0.137 | 0.0570 | 210 | <0.001 | 0.552 | 193 | Quadratic | -0.0360 ([ML] - 24.9)<sup>2</sup> + 128 |
| with O & S | 0.312 | 0.00300 | 243 | 0.00119 | 0.335 | 233 | Quadratic | -0.0484 ([ML] - 23.8)<sup>2</sup> + 128 |
| Lignin remaining |  |  |  |  |  |  |  |  |
| Single | <0.001 | 0.804 | 67.5 | 0.701 | 0.778 | 69.2 | Linear | -0.668 [ML] + 100 |
| with O | <0.001 | 0.916 | 61.1 | 0.0262 | 0.960 | 55.1 | Linear | -0.782 [ML] + 102 |
| with S | <0.001 | 0.864 | 68.5 | 0.167 | 0.888 | 67.4 | Linear | -0.899 [ML] + 105 |
| with O & S | 0.00142 | 0.811 | 60.6 | 0.0150 | 0.938 | 52.3 | Linear | -0.746 [ML] + 104 |
| Sasa (S) |  |  |  |  |  |  |  |  |
| Carbon remaining |  |  |  |  |  |  |  |  |
| Single | <0.001 | 0.962 | 126 | 0.0144 | 0.970 | 121 | Linear | -1.12 [ML] + 98.1 |
| with O | <0.001 | 0.969 | 132 | 0.954 | 0.967 | 134 | Linear | -1.13 [ML] + 101 |
| with B | <0.001 | 0.944 | 134 | 0.875 | 0.941 | 136 | Linear | -1.14 [ML] + 98.9 |
| with O & B | <0.001 | 0.943 | 150 | 0.729 | 0.941 | 152 | Linear | -1.46 [ML] + 101 |
| Nitrogen remaining |  |  |  |  |  |  |  |  |
| Single | 0.236 | 0.021 | 231 | 0.302 | 0.0260 | 232 | None | None |
| with O | 0.529 | -0.026 | 235 | 0.0486 | 0.110 | 232 | None | None |
| with B | 0.0287 | 0.163 | 223 | 0.261 | 0.175 | 223 | None | None |
| with O & B | 0.604 | -0.029 | 254 | 0.723 | -0.0660 | 256 | None | None |
| Lignin remaining |  |  |  |  |  |  |  |  |
| Single | 0.961 | -0.142 | 75.7 | 0.00698 | 0.639 | 66.0 | Quadratic | -0.0524 ([ML] - 20.6)<sup>2</sup> + 124 |
| with O | 0.703 | -0.118 | 73.2 | 0.0368 | 0.405 | 68.1 | Quadratic | -0.0579 ([ML] - 18.8)<sup>2</sup> + 121 |
| with B | 0.992 | -0.143 | 81.1 | <0.001 | 0.983 | 43.8 | Quadratic | -0.0933 ([ML] - 18.9)<sup>2</sup> + 134 |
| with O & B | 0.298 | 0.041 | 77.0 | 0.270 | 0.119 | 76.9 | None | None |

